# Bacterial population-level trade-offs between drought tolerance and resource acquisition traits impact decomposition

**DOI:** 10.1101/2024.06.22.600187

**Authors:** Ashish A. Malik, Jennifer B. H. Martiny, Antonio Ribeiro, Paul O. Sheridan, Claudia Weihe, Eoin L. Brodie, Steven D. Allison

**Affiliations:** School of GeoSciences, University of Edinburgh, Edinburgh, UK; School of Biological Sciences, University of Aberdeen, Aberdeen, UK; Department of Ecology & Evolutionary Biology, University of California, Irvine, USA; Centre for Genome-Enabled Biology and Medicine, University of Aberdeen, Aberdeen, UK; School of Biological and Chemical Sciences, University of Galway, Ireland; Earth and Environmental Sciences, Lawrence Berkeley National Laboratory, Berkeley, CA; Department of Environmental Science, Policy and Management, University of California, Berkeley, USA; Department of Earth System Science, University of California, Irvine, USA

## Abstract

Microbes drive fundamental ecosystem processes such as decomposition. Environmental stressors are known to affect microbes, their fitness, and the ecosystem functions that they perform, yet understanding the causal mechanisms behind this influence has been difficult. We used leaf litter on soil surface as a model in situ system to assess changes in bacterial genomic traits and decomposition rates over 18 months with drought as a stressor. We hypothesized that genome-scale trade-offs due to investment in stress tolerance traits under drought reduce the capacity for bacterial populations to carry out decomposition; and that these population-level trade-offs scale up to impact emergent community traits thereby reducing decomposition rates. We observed drought tolerance mechanisms that were heightened in bacterial populations under drought, identified as higher gene copy numbers in metagenome-assembled genomes. A subset of populations under drought had reduced carbohydrate-active enzyme genes which suggested – as a trade-off – a decline in decomposition capabilities. These trade-offs were taxonomically structured as distinct patterns appeared in populations across families. We show that trait-tradeoffs in bacterial populations under drought could scale up to reduce overall decomposition capabilities and litter decay rates. Using a trait-based approach to assess the population ecology of soil bacteria, we demonstrate genome-level trade-offs in response to drought with consequences for decomposition rates.

## Introduction

Modern omics technologies enable taxonomic and functional characterisation of microbiomes to shed new light on how microbes respond to change (1). These tools have transformed microbial ecology by providing a granularity needed to study complex and highly diverse environmental microbiomes (2). However, linking microbiome characteristics to ecosystem processes or to changes in pools and fluxes has remained difficult (3). This limitation may be in part because of methodological challenges in testing how environmental factors affect individual- or population-level physiology which ultimately impacts microbial fitness and contribution to ecosystem functioning. For instance, shotgun sequencing approaches often produce community-aggregated measurements that represent the additive properties of phylogenetically diverse members of the community, which may not reflect emergent properties resulting from complex interactions among community members and their environment (4, 5). Thus, the measured microbial and environmental characteristics may be decoupled. Therefore, there is a need to study microbial physiology at multiple levels of biological organisation – from populations of a species/strain to collective communities – to better understand and predict the ecosystem-level impacts of microbial processes.

Environmental stress can change cellular resource allocation with consequences for ecosystem processes. However, determining the quantitative impacts of environmental stressors on the metabolic response in distinct populations is extremely challenging. A trait-based approach, akin to that widely used in plant ecology (6, 7), offers the opportunity to functionally characterise and represent the enormous diversity of microbes involved in system-level processes such as decomposition. Traits are an organism ‘s phenotypic characteristics that govern process rates (5, 8). By focussing on the phenotypic aspects of microbes rather than their taxonomic identity, trait-based approaches provide a way forward to integrate functional information across species, space, and time. Quantitative phenotypic measurements have been made successfully at the community level, for example of carbon use efficiency (9) but are extremely challenging in single microbial populations in environments such as soil. However, we can use genetic markers to study traits in populations of single species or strains by leveraging new genome assembly and binning approaches that make it possible to extract hundreds of microbial genomes (metagenome-assembled genomes or MAGs) from environmental microbiomes (10, 11).

Traits can trade-off due to adaptive physiological processes in response to short-term fluctuations in the environment. Physiological constraints in stressful conditions such as reduced water availability could lead to greater cellular-level allocation to maintenance and survival relative to resource acquisition traits (12). If the environmental stressors persist over longer time periods, these will manifest as genome-encoded trade-offs across populations through ecological selection or evolutionary processes (13, 14). Reduced resource acquisition traits at the community level can have system-level impacts such as reduction in decomposition rates. However, such ecosystem-level quantifications or predictions are hard to make due to mismatches in functional response at different levels of biological organisation. Organismal level trade-offs in traits can be detected at the level of populations. Various ecological and abiotic factors may impact the emergent community response, potentially reinforcing or attenuating the population-level trade-offs in traits.

Climatic extremes such as drought are becoming more frequent and severe, and act as stressors for microbial decomposers. Drier environments are thought to reduce microbial activity and therefore organic matter decomposition (15, 16). Drought-induced reduction in microbial activity may occur due to organismal responses to water stress as well as due to limitations on resource diffusion and transport (15, 16). Drought is known to change the composition of active members of the microbial community through environmental filtering, as selective pressures enable some taxa to gain competitive advantage over others (17–19). Microorganisms also have the potential to acquire new genes through horizontal gene transfer or homologous recombination (20). Such evolutionary processes could enable organisms under chronic drought to gain stress tolerance traits. This functional diversification through generation of new genetic variation without shifting community taxonomic composition highlights why linking of microbial and ecosystem processes could be better achieved through trait-based approaches.

While population-level analysis helps reveal the organismal response to stress, the emergent community-level analysis is the most relevant to the impact of microbial processes at the ecosystem scale. The effect of stress on genomic traits in microbial populations and trade-offs with fitness traits should scale up to the community-level with consequences for ecosystem-level process rates. We used long-term drought as a persistent stress on microbes that grow on plant leaf litter to investigate the impact of stress on microbial traits at the population level and its impact on emergent community traits that influences rates of litter decomposition. We hypothesized that drought imposes constraints on the metabolism of decomposer populations such that increased investment in stress tolerance traits reduces resource acquisition traits. This trade-off scales up to reduce the emergent community-level decomposition capabilities and decrease rates of organic matter decomposition.

To test this hypothesis, we measured traits in populations of single species or strains and collective communities in an *in-situ* decomposition experiment in Mediterranean grass and shrub ecosystems. We used two different litter types to assess if microbial drought response strategies differ across litter of divergent chemical quality. The study was performed in litter bags of 1mm mesh size that were placed on the soil surface in experimental plots with continued drought treatment. Genomic traits of the decomposer community in litter bags were measured using shotgun metagenomics at four time points over an 18-month period (Figure S1). We used bacterial MAGs to represents populations of single species or strain and probed for trade-offs in traits using the frequencies of genes linked to traits of interest. The abundance of these populations in communities across treatments was used to compare trait trade-offs at the population and community levels. This rarely used approach tests if patterns across biological levels are scale-dependent and helps better understand the ecosystem-level implications of microbial decomposition.

## Results and discussion

### Ecosystem-level decomposition rates under drought

In contrast to our hypothesis, litter decomposition rates were unaffected by long-term drought (Figure 1; ANOVA p>0.05), although there was a non-significant trend towards lower decomposition rates in grass litter under reduced precipitation treatment compared to the ambient control. After 14 months of in situ experimental incubation in the grassland system, mass loss under reduced precipitation treatment was 50.5 ± 3.3 % compared to 53.8 ± 7.2 % under ambient precipitation treatment. Decomposition rates were significantly lower in shrub litter (ANOVA p<0.001) likely due to its higher C:N ratio, higher proportion of lignin and lower proportion of cellulose, hemicellulose and soluble compounds than the grass litter (21). After 14 months, mass loss in the shrubland system under reduced precipitation treatment was 41.8 ± 1.2 % compared to 37.3 ± 4.5 % under ambient precipitation treatment. Litter mass loss measures varied greatly across replicates due to soil infiltration into the litter bags, which could have affected the accuracy of the rate estimates (especially at T4, 18 months of incubation; this time point was excluded). Litter decomposition rates from other experiments performed at the same site using litter bags with smaller mesh size have found lower decomposition rates in grass litter under drought (22, 23). Although we did not find a statistically significant trend, drought can negatively impact decomposition in our experimental site as well as in other Mediterranean semi-arid ecosystems (24–27).

**Figure 1:**
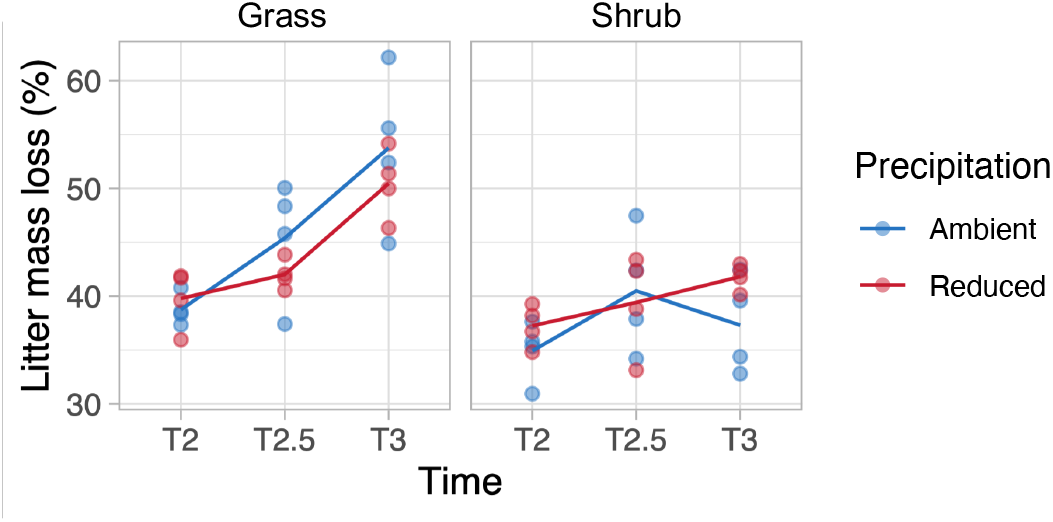
Effect of drought on decomposition rates. Decomposition rates measured as percentage loss of litter mass since the start of the litter bag experiment

*Community-level drought stress response linked to osmotic adaptations and iron metabolism* Metagenomic sequencing across the four sampled time points was used to investigate microbial community-level patterns in trait variation during decomposition under drought. We assessed changes in community-level functional gene abundance using metagenomic reads annotated with SEED Subsystems databases. Three categories of level 1 functions in Subsystems classification were significantly higher (p < 0.001) in the drought treatment relative to ambient: membrane transport, stress response, and iron acquisition and metabolism (Figure 2a). In microbial communities from both grass and shrub litter, sequence reads annotated to these three categories were consistently higher in drought treatment at most time points (Figure 2b) suggesting a similar drought response in very distinct communities (28, 29). Individual genes belonging to the three drought-enriched functional categories were linked to oxidative and osmotic stress, antiporters, ABC transporters, protein secretion systems and siderophores (Figure S2). Enrichment of these genes is consistent with our hypothesis and other studies of bacterial drought response pathways (28, 30, 31).

**Figure 2:**
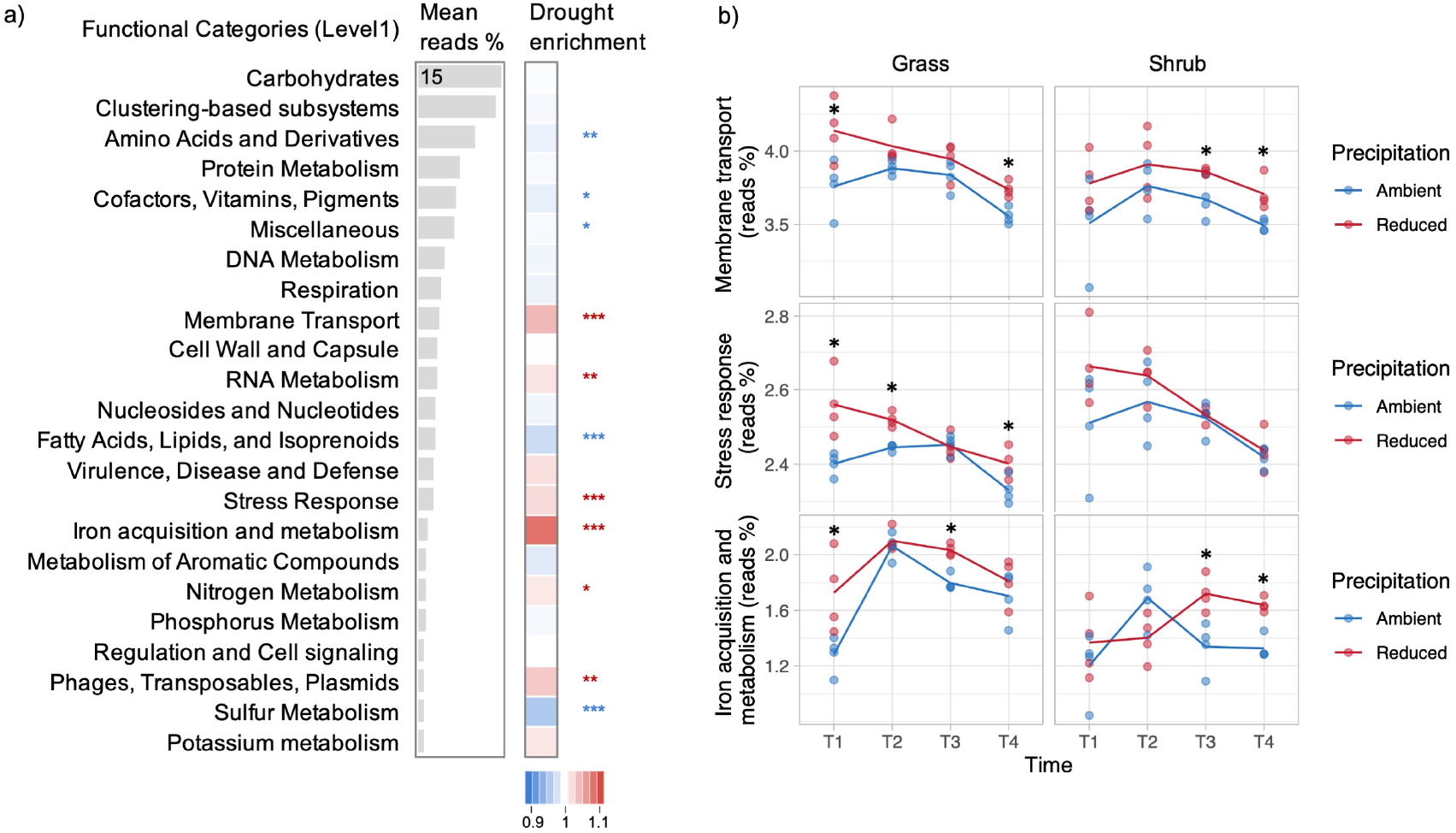
Community-level drought-enriched functional categories. **a**: Results of community-aggregated read abundance-based analysis of major categories of functional genes in grass and shrub litter across all time points. This differential abundance analysis highlighted the drought-enriched functional categories that were significantly higher in the reduced precipitation treatment compared to ambient. The bar plot shows mean read abundance and heatmap shows the fold change in response to drought. Asterisks indicate significant drought enrichment measured as pairwise variation between ambient and reduced precipitation treatments (*** p < 0.001, ** p < 0.01, * p < 0.05). **b**: Temporal patterns of community-aggregated read abundance across treatments for the most drought-responsive categories of functional genes: membrane transport, stress response, and iron acquisition and metabolism. Asterisks indicate significant pairwise difference between ambient and reduced precipitation treatments at each time point (* p < 0.05).

### Population-level drought stress response

To investigate drought responses in bacterial populations, we analyzed gene profiles in MAGs retrieved by de novo assembly and binning. Overall, we retrieved 533 bacterial MAGs with contamination of <10% and completeness of >45% (207 MAGs had completeness of >70%). These MAGs spanned all major bacterial phyla. In some cases, multiple MAGs were retrieved for a species which indicates strain-level genomic resolution; hence we consider MAGs representative of populations. For each MAG, we obtained gene copy numbers with KEGG annotations. From each of the three categories of functional genes that were most responsive to drought treatment at the community level (Figure 2), we chose the most abundant genes and queried for them in MAGs to assess whether they were present in higher copy numbers in populations under drought (gene copy numbers were normalized to account for variable genome size and bin completeness). We used the sum of copy numbers for the following set of genes and refer to them as drought responsive functions: (i) Na+:H+ antiporter in the ‘membrane transport ‘ category (Na+:H+ antiporter NhaA, NhaB and NhaC family; multicomponent Na+:H+ antiporter subunit A, B, C, D, E, F and G), (ii) glycine transport in the ‘stress response ‘ category (glycine betaine/proline transport system ATP-binding protein, permease protein and substrate-binding protein; choline/glycine/proline betaine transport protein; glycine betaine transporter; D-serine/D-alanine/glycine transporter) and (iii) Fe^3+^ transport in the ‘iron acquisition and metabolism ‘ category (ferric enterobactin transport system substrate-binding protein, permease protein and ATP-binding protein; ferric hydroxamate transport system substrate-binding protein and permease protein; ferric transport system ATP-binding protein, permease protein and substrate-binding protein).

For all three drought responsive functions, mean gene copy numbers were higher in bacterial MAGs from drought communities relative to control in both grass and shrub litter (Figure 3a-c). This difference suggests that bacterial populations under drought have increased gene copy numbers for key stress tolerance traits (Figure 3d-f) which likely improved their fitness under drought, consistent with our hypothesis. Membrane transport functions performed by Na+:H+ antiporter subunit A in maintaining monovalent cation and proton homeostasis could be crucial for drought stress tolerance, a mechanism that has been identified in microbes and plants to tolerate drought and salinity stress (32–34); we have previously reported this adaptation in our study at the same field site using metatranscriptomics (28). Differential abundance of genes for transport of osmolytes such as glycine betaine and proline under osmotic stress has been well documented in laboratory cultures and to some extent in soils (35, 36). These genes are related to maintenance of intracellular osmotic potential by increasing the concentrations of organic compatible solutes. In addition to osmotic stress, drought appears to reduce iron availability in microorganisms due to reduced diffusion and increased aeration that oxidizes Fe^2+^ into insoluble Fe^3+^. We demonstrate higher abundance of genes for iron transport and metabolism (e.g., Fe^3+^ siderophore transport system) presumably as a metabolic adaptation to mitigate iron limitation caused by drought (37–39).

**Figure 3:**
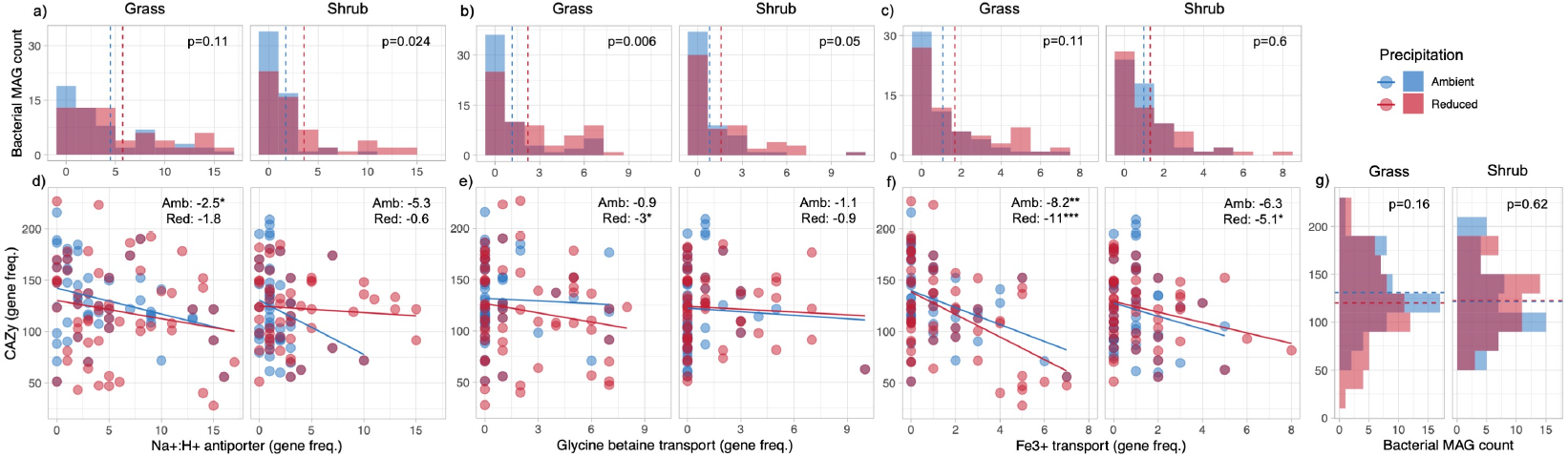
Gene copy numbers of drought responsive functions within MAGs and their population-level trade-offs with carbohydrate-active enzyme (CAZy) genes. **a-c:** Histograms showing the count of all bacterial metagenome-assembled genomes (MAGs) with varying number of genes for the drought-enriched functions: Na+:H+ antiporter (a), glycine betaine (b), and iron transport (c). The vertical dotted lines represent mean number of genes in MAGs from ambient (blue) and reduced (red) precipitation treatment. Also presented are the p values of Wilcoxon signed rank sum test to highlight statistical differences in number of genes across the precipitation treatments. **d-f**: Scatter plot of number of genes for drought-enriched functions in each MAG on the x-axes and total CAZy genes on the y-axes with linear regression lines for ambient (blue) and reduced (red) precipitation treatment. Also presented are the slope values for each regression and asterisks which indicate significant negative correlation of drought enriched genes and CAZy genes (*** p < 0.001, ** p < 0.01, * p < 0.05). **g**: Histogram showing the count of bacterial MAGs with varying number of CAZy genes. The horizontal dotted lines represent mean number of genes in MAGs from ambient (blue) and reduced (red) precipitation treatment and statistical differences are highlighted using p values of the Wilcoxon signed rank sum test.

### Population-level trade-offs between stress tolerance and resource acquisition traits

We tested whether increased investment in stress tolerance traits in populations under drought (measured here in terms of higher gene copy numbers for drought responses) leads to lower decomposition capabilities. To demonstrate such a tradeoff, we quantified the total number of CAZyme (carbohydrate-active enzyme) genes in the recovered MAGs; CAZyme genes are involved in decomposition of carbohydrates (40), a major substrate in grass and shrub litter. A trade-off between drought stress tolerance and resources acquisition traits would manifest in a negative correlation between counts of genes representing the two traits, with the slope determining the magnitude of this trade-off. We expected to see this trade-off in populations under both drought and control treatments, suggesting that the trade-off may occur widely in populations as a longer-term evolutionary response (13, 14) but we anticipated that the magnitude of that trade-off would be higher under drought due to selection of drought tolerant populations.

We observed statistically significant negative relationships in grass bacterial MAGs, and the slopes were either similar or more negative in the drought treatment compared to control (Figure 3d-f). In other words, populations in grass litter with higher gene copy numbers for the drought responsive functions had lower total CAZyme genes, consistent with our trade-off hypothesis. This pattern was also seen in bacterial populations from shrub litter (Figure 3d-f), but it was weaker compared to populations from grass litter. Interestingly, the negative relationship between Fe^3+^ transport genes and total CAZyme genes was stronger and statistically significant in drought treatments from both grass and shrub litter, likely because ‘iron transport and metabolism ‘ was the most highly enriched functional gene category in response to drought in both grass and shrub communities (Figure 2). We also observed that a higher number of bacterial MAGs in grass litter under drought compared to ambient had a lower count of CAZyme genes (Figure 3g). Such a drought-linked pattern was not observed in bacterial MAGs from shrub litter (Figure 3g). The negative relationship between drought-enriched genes and total CAZyme genes demonstrates the presence of a gene-level tradeoff between stress tolerance and resource acquisition traits in bacterial populations, a novel finding in microbial ecology.

### Population-level trade-offs were taxonomically structured

To further probe these novel population-level trade-offs, we examined the influence of reduced precipitation on the abundance of bacterial populations over the 18-month decomposition experiment. Abundance of populations (individual MAGs) was quantified as genome copies per million metagenomic reads across all samples. Instead of analyzing all populations that were present in the precipitation treatments as in Figure 3, we identified drought-selected populations as MAGs that were significantly more abundant in drought samples compared to the ambient control and vice versa. We obtained the following number of MAGs that were differentially abundant in each treatment: grass ambient precipitation: 25, grass reduced precipitation: 23, shrub ambient precipitation: 9, shrub reduced precipitation: 18 (low abundance MAGs with less than 5 genome copies per million reads were excluded). For each MAG, we used the sum of copy numbers for the genes representing all three drought responsive functions as a measure of drought stress tolerance and plotted it against total CAZyme genes (Figure 4a) while also accounting for the taxonomic affiliation of each MAG at the level of family (coarser level when this was not available). We demonstrate that population level trade-offs were only observed in grass litter under drought and that these trade-offs were taxonomically structured.

**Figure 4:**
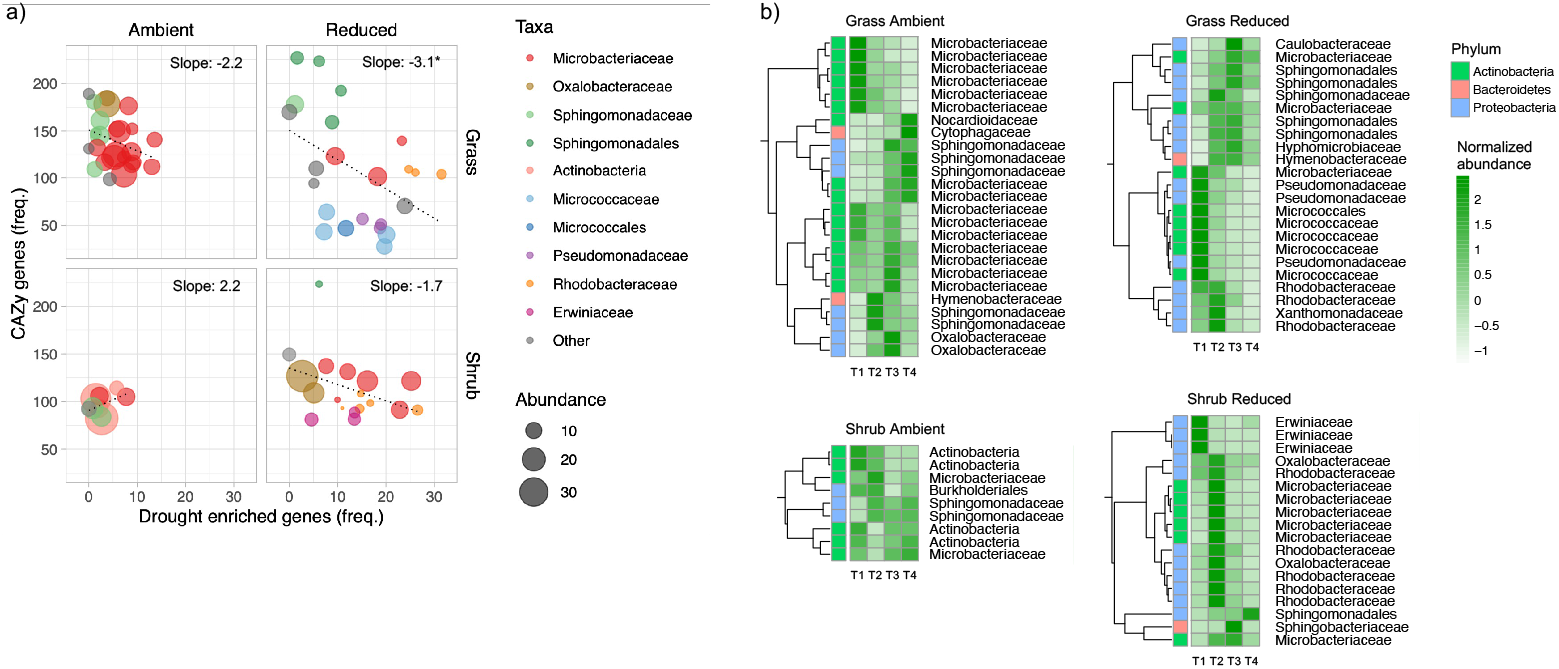
Bacterial population-level trade-offs are driven by taxonomic differences. **a**: Scatter plot of sum number of genes for all drought-enriched functions in MAGs on the x-axes and total CAZy genes on the y-axes. Displayed here are only those MAGs that were differentially abundant across the precipitation treatments. Each point is a bacterial MAG, and its size represents its mean abundance in metagenomes across all four time points expressed as genome copies per million reads. Linear regression lines and the slope values for each regression are presented. Very low abundance MAGs (<5 genome copies per million reads for bacterial MAGs) were excluded from this analysis. The color represents the taxonomic affiliation of MAGs at the level of family (or order/phylum when that was not available). **b**: Heatmaps showing the abundance of the same MAGs at individual time points across all treatments. Similar temporal patterns are clustered together, and the color codes represent their phylum level affiliation.

Drought-selected populations in grass litter belonging to the taxa Micrococcaceae/Micrococcales (Actinobacteria) and Pseudomonadaceae (Proteobacteria) have reduced resource acquisition traits with a below average (<100 genes) CAZyme count (Figure 4a). However, there were other drought-selected populations belonging to the taxa Microbacteriaceae (Actinobacteria), Sphingomonadaceae or Sphingomonadales (Proteobacteria) and Rhodobacteraceae (Proteobacteria) that maintained a higher CAZyme gene count (>100) while also having higher drought enriched genes (Figure 4a). This demonstrates that the population level trade-offs observed in grass litter under drought were driven by taxonomic difference at the level of family; some of the families identified have been extensively studied at this field site (41, 42). We then analyzed the temporal dynamics in the abundance of these populations (Figure 4b) and found that the drought-selected populations with a lower CAZyme count were more abundant at the start of the decomposition process (T1). In contrast, the drought-selected populations with a higher CAZyme count were more abundant at the later time points (T2 to T4). Drought-selected populations in shrub litter did not demonstrate a reduction in CAZyme gene count (Figure 4a). However, populations belonging to the family Erwiniaceae (Proteobacteria) that were more abundant at the start of the decomposition process (Figure 4b) had below average CAZyme count (Figure 4a) reinforcing that trade-offs were driven by family-level taxonomic differences. Interestingly though, trait distributions were quite varied in populations within the family Microbacteriaceae which suggests that trade-offs may also occur across populations within a family.

### Linking population-level traits to decomposition rates

The results from our 18-month decomposition experiment suggest that drought selects for bacterial populations with gene-level stress tolerance traits that could confer a competitive advantage. We show that in some drought-selected populations, enhanced stress tolerance strategies negatively impacted decomposition capabilities as inferred from CAZyme gene counts. We also observed that the drought-related trade-offs appear across families, and potentially also across populations within families. Our genome-based analysis highlights gene-encoded patterns that likely appear as drought-adapted bacterial populations are favored by the decade-long drought imposed in our experiment. These population-level trade-offs in traits in response to long-term drought were observed in grass litter and were weak or absent in shrub litter. Tradeoffs between stress tolerance and resource acquisition traits in bacterial populations could reduce decomposition capabilities and ecosystem-level litter decay rates under drought. Organismal response of bacteria to drought in terms of physiological changes is one of the factors that affects decomposition rates. It is likely that other factors such as changes in fungal traits, limitations to resource diffusion and transport, changes in plant litter chemistry and physico-chemical factors also affect ecosystem-level decomposition rates (16, 43) thereby decoupling microbial decomposer traits at the population level from ecosystem process rates. Nevertheless, we show clear links between the metabolism of bacterial populations and collective emergent traits. When integrated into a biogeochemical framework (5), such trait-based scaling can be applied to link the metabolism of microbial populations in a changing environment to the ecosystem functions they perform.

## Materials and Methods

### Field site

We performed a leaf litter decomposition experiment in the field at the Loma Ridge Global Change Experiment situated near Irvine, California, USA (33°44′N, 117°42′E, 365 m elevation). The site experiences a Mediterranean climate (mean annual temperature: 17°C, mean annual precipitation: 325 mm) with a summer drought from May to October and periods of precipitation from November to April. The vegetation at the site consists of an annual grassland adjacent to a coastal sage scrub ecosystem (22, 44). The long-term experiment consisted of grassland plots (6.7 × 9.3 m) with native perennial grass *Stipa pulchra*; exotic annual grasses such as *Avena, Bromus, Festuca*, and *Lolium*; and forbs such as *Erodium* and *Lupinus*. The shrubland plots (18.3 × 12.2 m) host crown-sprouting shrub species such as *Salvia mellifera, Artemisia californica, Eriogonum fasciculatum, Acmispon glaber* and *Malosma laurina* (29, 44). We used the long-term drought experimental plots that received continuous field precipitation manipulations since 2007; reduced precipitation treatment plots were covered with retractable clear polyethylene rain shelters during a subset of precipitation events during the wet season to achieve ∼40 % precipitation reduction compared to ambient plots (Figure S1). We used a total of 16 plots consisting of four replicated plots per treatment of grassland ambient, grassland reduced, shrubland ambient and shrubland reduced.

### Experimental design

Litter of four types was collected from each treatment on 30 August 2017. Litter from all replicated plots (n=4) within each treatment was homogenised by hand mixing. To make litter bags, dry, senescing sheath and blade material in grassland was cut to a length of approximately 10cm, whereas freshly fallen intact dry leaves were used in shrubland. 6 g dry litter mass was placed into 15 cm × 15 cm bags of 1 mm mesh window screen. Litter bags were deployed in the field on 12 September 2017; they were placed on top of the soil surface under the canopy. 16 litter bags were collected at each time point (Figure S1): 30 November 2017 (T1; end of the dry season), 11 April 2018 (T2; end of the wet season), 16 July 2018 (T2.5; middle of the dry season; additional sampling point only to measure litter mass loss), 5 November 2018 (T3; end of the dry season) and 19 February 2019 (T4; end of the wet season). Litter in each bag was weighed at the start of the experiment and at each sampling point. A subsample was dried to constant mass at 65°C to obtain the moisture content and dry mass of litter. Mass loss is reported as percentage initial dry mass.

### DNA extraction and sequencing

DNA was extracted from a coarse-ground litter aliquot of 50 mg for time points T1, T2, T3 and T4 (total samples: 64). We used ZymoBiomics DNA isolations kits (Zymo Research, Irvine, CA, USA) and followed manufacturer instructions. Sample was homogenized by bead beating for 5 min at maximum speed (6.0 m/s, FastPrep-24 High Speed Homogenizer, MP Biomedicals, Irvine, CA, USA). Purity and concentration of extracted DNA was assessed using gel electrophoresis, a Qubit fluorometer (LifeTechnologies, Carlsbad, CA, USA) and Nanodrop 2000 Spectrophotometer (Thermo Scientific, USA). Metagenomics library preparation and sequencing were carried out at the DNA Technologies and Expression Analysis Cores at the University of California Davis Genome Center. We used PE150 sequencing on NovoSeq platform (Illumina, San Diego, CA, USA) with the default insert size of 250-400bp.

### Reads-based analysis to obtain community-level functional gene abundances

To get a reads-based assessment at the community-level, DNA sequences were annotated with the Metagenomics Rapid Annotation using Subsystems Technology (MG-RAST) server version 4.0.3 (45). Functional annotations were performed with the SEED Subsystems database and taxonomic classification up to genus level was performed using the RefSeq database (maximum e-value cut-off of 10^−5^, minimum identity cut-off of 60% and minimum length of sequence alignment of 15 nucleotides).

### Metagenomic co-assembly, binning and annotation of prokaryotic genomes

For population-level analysis of genetic traits, the goal was to retrieve a high number of metagenome-assembled genomes (MAGs) by de novo assembly and binning. To achieve this, we used a co-assembly approach merging the four replicates per treatment. Metagenome Orchestra-MAGO (version V2.2b; 2020-03-08) (46) was used to produce MEGAHIT (version 1.2.8) (47) co-assemblies. Quality control and trimming approaches were like those used for individual assemblies. To extract prokaryotic bins from co-assemblies, we used the binning module of MetaWRAP (version 1.2.1) (48), MetaBAT2 (version 2.12.1) (49), Concoct (version 1.0.0) (50), and Maxbin2 (version 2.2.6) (51). MetaWRAP ‘s downstream modules were run over the collected bins with thresholds of minimum 45% completion and maximum 10% contamination obtaining bin-related statistics using CheckM (version 1.0.12) (52). For the resulting bins, carbohydrate active enzymes (CAZy) were annotated using dbCAN2 (version 2.0.11) (53) with default parameter values. To obtain KEGG orthology (KO) annotations for the bins, we used Prodigal (version 2.6.3) (54) to carry out gene-calling over the bins and output was then used to query the KOfam database (ftp://ftp.genome.jp/pub/db/kofam/) with kofamscan (version 1.2.0) (55) using default parameter values. The quant_bins module of MetaWRAP (version 1.2.1) was used to compute the abundances of bins across samples.

### Statistical analysis and visualisations

Statistical significance of decomposition rates measured as litter mass loss was estimated across vegetation and precipitation treatments using multiple ANOVA. Categories of functional genes (level 1 of SEED Subsystems classification) with the highest and most statistically significant fold changes in read abundance in reduced precipitation treatment compared to the ambient were identified as drought-enriched functions. Fold change was estimated as the ratio of sum-normalised read abundance in reduced and ambient precipitation treatments averaged across all time points and vegetation types. Statistical significance of this change with precipitation was estimated using one-way ANOVA across the precipitation treatments. Three functional categories with p < 0.001 were identified: membrane transport, stress response, and iron acquisition and metabolism. We then used differential abundance analysis carried out using DESeq2 (56) with precipitation as the experimental condition to identify the individual drought-enriched functional genes within the three categories. The temporal pattern and treatment effect of the three drought-enriched functional groups was visualised using line plots. Statistical significance of change with precipitation treatment at each time point was estimated using one-way ANOVA. Temporal pattern and treatment effect of individual drought-enriched functional genes were visualised using a heatmap created with the pheatmap package (57); more descriptive level 2 functional annotation of each gene was highlighted using a colour code.

To demonstrate gene level trade-offs between drought-enriched genes and genes for decomposition, we made scatter plots with the sum of normalized copy numbers for multiple genes representing the three abundant drought responsive gene functions on the x-axis and normalized total number of CAZy genes on the y-axis. Normalization was performed using the total gene number per genome which accounted for genome size and bin completeness. Trade-offs were visualized as negative regressions, displayed on the scatter plot using regression lines. Bacterial MAG-level gene copy numbers for total CAZy genes and drought enriched genes across treatments were visualised using geom_point and geom_smooth lm function. Bacterial MAG count for variable gene copy numbers for total CAZy genes and drought enriched genes were plotted with geom_histogram; geom_vline function was used to indicate the mean gene copy number across all MAGs. Wilcoxon signed rank sum test was used to examine the statistical differences in gene copy number across the precipitation treatments. The taxonomic structuring of the trade-offs in resource acquisition and stress tolerance traits was plotted with geom_point and geom_smooth lm function. Here, only a subset of the MAGs from each treatment were plotted; these were MAGs that were differentially abundant in drought samples compared to the ambient control and vice versa. Trade-offs visualized as negative regressions were displayed on the scatter plot using regression lines. Abundance of MAGs in metagenomes was obtained using the quant_bins function expressed as genome copies per million reads and visualised as the size of each point representing the mean abundance over the four time points. The temporal pattern of these differentially abundant MAGs across the different treatments were plotted as a heatmap with the pheatmap package; phylum-level affiliation of each MAG was highlighted using a colour code. MAGs were labelled as per their family-level affiliation; when this was not available a coarser taxonomic classification at the level of order or phylum was used. All visualisations were made in ggplot2 package (58) in R 3.4.2 (59).

## Author Contributions

AAM, JBHM, ELB and SDA designed research; AAM coordinated the project; AAM and CW were involved in litter sampling, experimental setup, sample processing and DNA extractions; AAM, AR and POS performed bioinformatic analyses; AAM conducted all the statistical analysis, JBHM, ELB and SDA contributed reagents and analytical tools; AAM drafted the manuscript with inputs from JBHM and SDA, and all authors were involved in critical revision and approval of the final version.

## Acknowledgments

We acknowledge funding from the US Department of Energy (DOE) Genomic Science Program, BER, Office of Science projects DE-SC0016410 and DE-SC0020382 awarded to University of California Irvine, and DE-AC02-05CH11231 awarded to Lawrence Berkeley National Laboratory. DNA sequencing was carried out at the DNA Technologies and Expression Analysis Cores at the UC Davis Genome Center, supported by NIH Shared Instrumentation Grant 1S10OD010786-01. Bioinformatics support was available through the Centre for Genome Enabled Biology and Medicine of the University of Aberdeen through an internal Research Enhancement Scheme grant.

