## Supplementary information for "Bacterial population-level trade-offs between drought tolerance and resource acquisition traits impact decomposition"

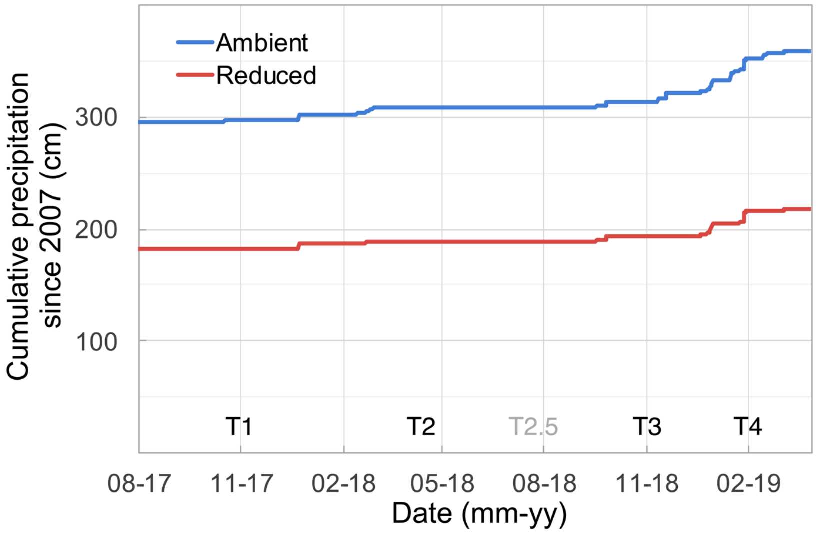


**Figure S1: Precipitation treatments and sampling points**. Cumulative precipitation since 2007 (start of the long-term drought experiment) in the ambient and reduced precipitation plots. Litter was collected in August 2017 from the four vegetation-precipitation treatment plots (grass ambient, grass reduced, shrub ambient, shrub reduced). Litter bags were deployed in the field plots in September 2017 and collected at four time points over 18 months which are labeled as T1-4; T2.5 was an additional sampling point only to measure litter mass loss.

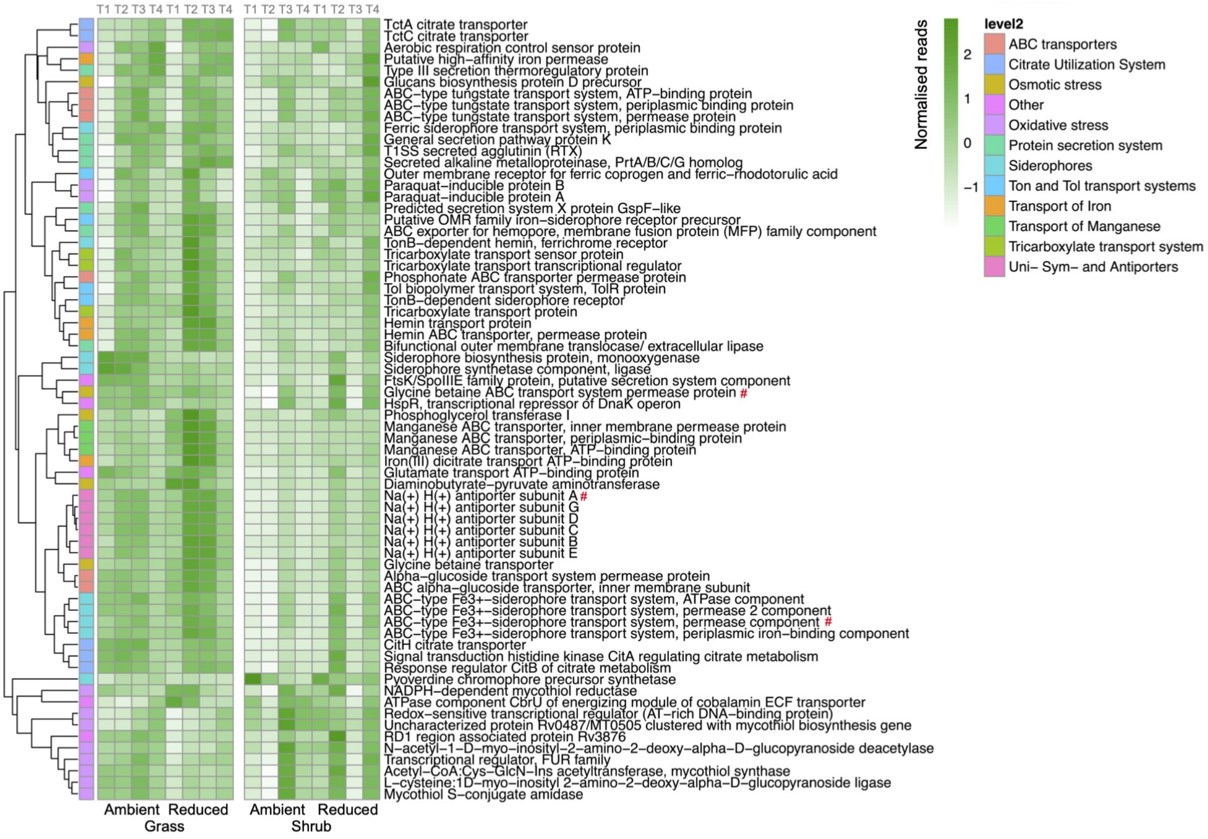


**Figure S2: Community-level distribution of drought-enriched genes**. Heatmap of temporal patterns of individual gene level differences in normalized read abundances across treatments for the most abundant functional genes in each of the three level 1 functional categories identified in Figure 2. Similar patterns across treatments are clustered together, and the color codes represent more descriptive level 2 categories of the functional genes. Red # highlights the most abundant drought-enriched functional genes in each of the three categories: Na+:H+ antiporter, glycine betaine transport system, and Fe3+ siderophore transport system.
